# Proteomic analysis revealed that the oomyceticide phosphite exhibits multi-modal action in an oomycete pathosystem

**DOI:** 10.1101/2022.08.16.504066

**Authors:** Christina E. Andronis, Silke Jacques, Francisco J. Lopez-Ruiz, Richard Lipscombe, Kar-Chun Tan

## Abstract

**BACKGROUND:** Phytopathogenic oomycetes constitute some of the most devastating plant pathogens that cause significant crop and horticultural loss. *Phytophthora cinnamomi* is a phytopathogenic oomycete that causes dieback disease in native vegetation and a variety of crops. This pathogen can survive through harsh environmental conditions which gives it an advantage over its susceptible hosts. The only implemented chemical used to control *P. cinnamomi* is the oomyceticide phosphite. Despite its widespread use, the mode of action of phosphite is not well understood and it is unclear whether it works directly on the pathogen or through the host. Additionally, resistance to phosphite is emerging in *P. cinnamomi* isolates and other oomycete phytopathogens.

**RESULTS:** The mode of action of phosphite on the pathogen and through a model host was investigated using label-free quantitative proteomics. *In vitro* treatment of *P. cinnamomi* with phosphite hinders growth by interfering with metabolism, signalling and gene expression, traits that are not observed in the tolerant isolate. When the model host *L. angustifolius* was treated with phosphite, enrichment of proteins that are associated with photosynthesis, carbon fixation and lipid metabolism in the host was observed. An increase in the production of a range of defence-related proteins was observed.

**CONCLUSION:** We hypothesise direct and indirect models of the multi-modal action of phosphite that directly targets the pathogen as well as alters plant metabolism and immune response.

## 1 INTRODUCTION

Phytopathogenic oomycetes are significant plant pathogens in natural ecosystems and agriculture. These pathogens cause substantial environmental and economic losses from plant death and management costs. *Phytophthora cinnamomi* is an oomycete that causes dieback disease in native vegetation and several crops including avocado, macadamia, pineapple, and a variety of stone fruits. This necrotic pathogen attacks the roots of susceptible hosts and causes plant death via saprophytic growth ^1^. The dynamic lifecycle of this organism drives its resilience and success as a plant pathogen ^2^. Similarly to other *Phytophthora spp*, its spores enable it to survive through harsh environmental conditions and thrive once conditions become more favourable. This gives the pathogen an advantage over its susceptible hosts. Characteristically, oomycetes produce an assortment of virulence molecules throughout their life cycles which facilitate the infection process ^3,4^.

Phosphite is the only commercially available chemical used to control *P. cinnamomi* ^5^. Inorganic phosphites are reduced forms of phosphate that are commonly used as oomyceticides to manage diseases caused by the *Phytophthora spp*. This includes *P. cinnamomi, P. nicotianae, P. palmivora, P. capsici* and *P. infestans*, along with other oomycetes such as downy mildews such as *Pseudoperonospora humuli* and *Bremia lactucae* ^6–10^. It is applied on native trees and horticultural crops by foliar spray or direct injection into the trunk where it circulates throughout the plant system ^11^. Phosphite cannot eradicate *P. cinnamomi* in the field but it can be used to reduce the severity and spread of the disease. It is also used as a preventative measure in uninfected areas ^11–13^. The only other strategies implemented to combat dieback disease are the use of tolerant root stocks if they are available (such as commercially available avocado root stocks) and hygiene measures applied to vehicles and personnel, which aim to minimise spread ^14–17^. *In vitro* fungicide screening of *P. cinnamomi* has shown growth inhibitory effects of chemical alternatives such as metalaxyl, fosetyl-A1, benzethonium chloride and copper salts however, these remain to be tested in the field ^18,19^. As such, phosphite remains the only option for the chemical control of *P. cinnamomi*.

Despite its widespread use in dieback management, the mode of action of phosphite is not well understood. It is suggested that it acts both directly on the pathogen and indirectly through the plant host by priming the plant immune system ^7,8,20–22^. It has been reported that mycelial growth and sporulation are inhibited by phosphite *in vitro* ^20,21^. *In planta, t*he increase in transcription of defence-related genes associated with the salicylic and jasmonic acid pathways in *Arabidopsis thaliana* and potato crops have been demonstrated post-treatment with phosphite ^7,8,22^. The effect of phosphite on potato leaves indicates an increase in defence responses and altered metabolism, glycolysis, and carbon fixation ^22^. These findings have led to the hypothesis that phosphite primes the immune system of plants for potential infection. Despite these observations, the mode of action of phosphite on the pathogen has not been defined, nor have the mechanisms of phosphite primed plant defence.

Tolerance of *P. cinnamomi* to phosphite is widespread with most reports originating from horticultural plantations and variability in phosphite sensitivity has been demonstrated *in vitro* and *in planta* ^20,23^. Tolerance to phosphite has also been reported in *P. nicotianae, P. capsici, Bremia lactucae* and *Pseudoperonospora humuli* ^9,10,24–26^. This is a suspected result of prolonged use of phosphite in agriculture and poses a significant threat to natural ecosystems and the agricultural industry. These tolerant isolates are driving the pressure for improved management strategies.

To elucidate the mode of action of phosphite on the pathogen and the host, we used a label-free quantitative proteomic approach to conduct a study encompassing the effects of phosphite on a sensitive and a tolerant isolate of *P. cinnamomi*, as well as investigating the effects of phosphite on the physiology in a model plant system that is susceptible to dieback. *P. cinnamomi* is used as model to understand phosphite tolerance as it is treated with phosphite in the field and tolerant isolates have emerged. By using a shotgun approach, we can obtain a snapshot of the biochemical processes that are altered as a result of treatment with phosphite and determine whether phosphite exhibits a direct and/or indirect mode of action.

## 2 MATERIALS AND METHODS

### 2.1 *In vitro* treatment of *P. cinnamomi* with phosphite

Stocks of two *P. cinnamomi* isolates, MU94-48 and CPSM366 were obtained from the Centre of Phytophthora Science and Management, Perth, Australia. MU94-48 was collected from susceptible *Eucalyptus marginata* in Willowdale, Western Australia with no history of phosphite use and CPSM366 was collected from a Western Australian avocado orchard with a history of extensive phosphite use and reduced efficacy of protection on dieback was observed ^23,27^. To determine the level of phosphite sensitivity of both isolates, they were grown on Ribeiro’s media amended with phosphite at concentrations ranging from 0.0 μg mL^−1^ to 1000 μg mL^−1 28^. Mycelial cultures were incubated at room temperature in the dark. After 14 days of growth, the mycelial radial growth was measured to determine the growth inhibition. The EC_50_ and minimum inhibitory concentration (MIC) for both isolates were calculated as previously described ^29^. For proteomics analysis, growth inhibition of 0% and 20% were used as untreated and sub-lethal doses of phosphite to cause a physiological effect and provide enough biomass for experimentation. Mycelia were harvested for protein extraction by scraping from the surface of the plate, snap-frozen in liquid nitrogen and freeze-dried. Lyophilised mycelia were ground using metal beads and a tissue mill (Retsch, Haan, Germany) at a frequency of 3 Hz/s for three minutes.

### 2.2 *In planta* treatment of *Lupinus angustifolius* and inoculation with *P. cinnamomi*

The narrow leaf lupin *Lupinus angustifolius* (cv. ‘Tanjil’) seeds were surface sterilised with 5% sodium hypochlorite, washed twice with 70% ethanol and washed three times with water. Seeds were placed in clear containers lined with moist Whatman paper (Cytiva, Massachusetts, USA) and left to germinate for 3 days at room temperature under natural light ^30^.

Three-day-old germinated seedlings were sprayed with a 0.5% pH 7 solution of phosphite (Sigma, St Louis, USA) using a hand-triggered spray bottle and untreated seedlings were sprayed with water ^31–33^. Seedlings were incubated on a light shelf at room temperature with a 12-hour photoperiod and seedlings were sprayed daily with water ^30^. Lupin root tips were harvested one day post first treatment. 1.5 cm of root tips were excised and immediately snap frozen, freeze dry, and ground to a fine powder using metal beads and a tissue mill (Retsch, Haan, Germany) at a frequency of 3 Hz/s for three minutes. Three biological replicates (of which three root tips were collected) were used for each sample.

To test the effect of phosphite on the colonisation ability of *P. cinnamomi*, 24 hours post-spraying, the infected seedlings were inoculated with 5 mm mycelial discs of each isolate. Containers were placed back on the light shelf for three days and lesion scores from zero to three were recorded.

### 2.3 Protein extraction and digestion

300 μl of extraction buffer (25 mM Tris-HCl pH 7.5, 0.25% SDS, 50 mM Na_2_PO_4_, 1 mM Na_2_F, 50 μM Na_3_VO_4_, 1 mM phenylmethylsulfonyl fluoride and a protease inhibitor cocktail) was added to the ground mycelia and the samples were kept on ice for 30 minutes with regular gentle mixing. The crude extract was centrifuged at 20,000 g at 4 °C for 30 minutes, the solubilised proteins were decanted, and proteins were precipitated using six volumes of ice-cold acetone and incubated at −20 °C overnight ^34^.

300 μl of extraction buffer (125 mM Tris-HCl pH 7.0, 7 % SDS, 0.5 % PVP-40) was added to the whole ground plant material and the samples were kept on ice for 30 minutes with regular gentle mixing. The crude extract was centrifuged at 20,000 g at 4°C for 30 minutes. Proteins were purified by adding 800 μL of ice-cold methanol and 200 μL of ice-cold chloroform to 200 μL of the solubilised protein samples. Samples were vortexed, and 500 μL of water was added and centrifuged for 5 minutes at 15000g a 4 °C. The aqueous phase was removed and 500 μL of methanol was added. Samples were inverted and the supernatant was discarded. 1 mL of ice-cold acetone was added and the samples were incubated at −20 °C overnight ^35^.

The mycelial and root protein pellets were washed twice with ice-cold acetone and reconstituted in 200 μl of 0.5 M triethylammonium bicarbonate (pH 8.5) before reduction and alkylation with 20 μL of 50 mM tris(2-carboxyethyl)phosphine (Thermo Scientific, Massachusetts, USA) and 10 μL 200 mM methyl methanethiosulfonate. Samples were tryptically digested overnight at 37 °C at a ratio of 1:10, subsequently desalted on a Strata-X 33 um polymeric reverse phase column (Phenomenex, Torrance, CA, USA) and dried in a vacuum centrifuge ^36^.

### 2.4 Mass spectrometry and data analysis

1 μg of each sample was loaded on columns and peptides were resolved with a gradient of 10-40% acetonitrile (0.1% formic acid) at 300 nL/min over 90 minutes and eluted through a nanospray interface into a Q-Exactive Orbitrap mass spectrometer (Thermofisher Scientific). Qualitative and label-free quantification was performed using Proteome Discoverer 2.3 ^37^. Mass spectra from the *in vitro* assay were matched to the *P. cinnamomi* MU94-48 genome consisting of 26,151 protein-coding sequences ^38^. The proteomes between phosphite-treated MU94-48 and CPSM366 were compared to simultaneously gain insight into the effects of phosphite on sensitive *P. cinnamomi* and understand the differences between phosphite sensitivity and tolerance. The mass spectra obtained from the *in planta* assay were matched to the *L. angustifolius* genome consisting of 39,339 protein-coding sequences ^39^. For protein identification, 1 or more 95% confidence peptides were used. For label-free quantification, proteins with 2 or more 95% confidence peptides were used and for significant differential abundance a p-value threshold of <0.05 was used. Ratios and p values for quantitative analysis were generated using the default t-test hypothesis testing over biological replicates 37

To elucidate the functions of the detected proteins and understand the biochemical differentiation between samples, Gene Ontology (GO), KEGG and Interpro were used ^40–42^. For qualitative analysis, a Fisher’s exact test was performed on assigned GOs to indicate GO enrichment within samples compared to assigned ontologies of the whole genome annotation in each respective organismp.

## 3 RESULTS

### 3.1 Growth inhibition of *P. cinnamomi* isolates by phosphite

To determine the sensitivity of *P. cinnamomi* isolates MU94-48 and CPSM366 to phosphite, both isolates were grown on phosphite-supplemented media, and sensitivity was determined based on the radial growth of hyphal colonies. Both isolates exhibited a similar radial growth rate in the absence of phosphite (Figure 1). The MIC for the sensitive and tolerant isolates were 500 μg/ mL and >1000 μg/ mL respectively. At concentrations of 1000 μg/ mL the growth of CPSM366 was reduced but not completely inhibited. The EC_50_ for the MU94-48 and CPSM366 were 10.8 μg/ mL and 415.6 μg/ mL respectively.

**Figure 1.**
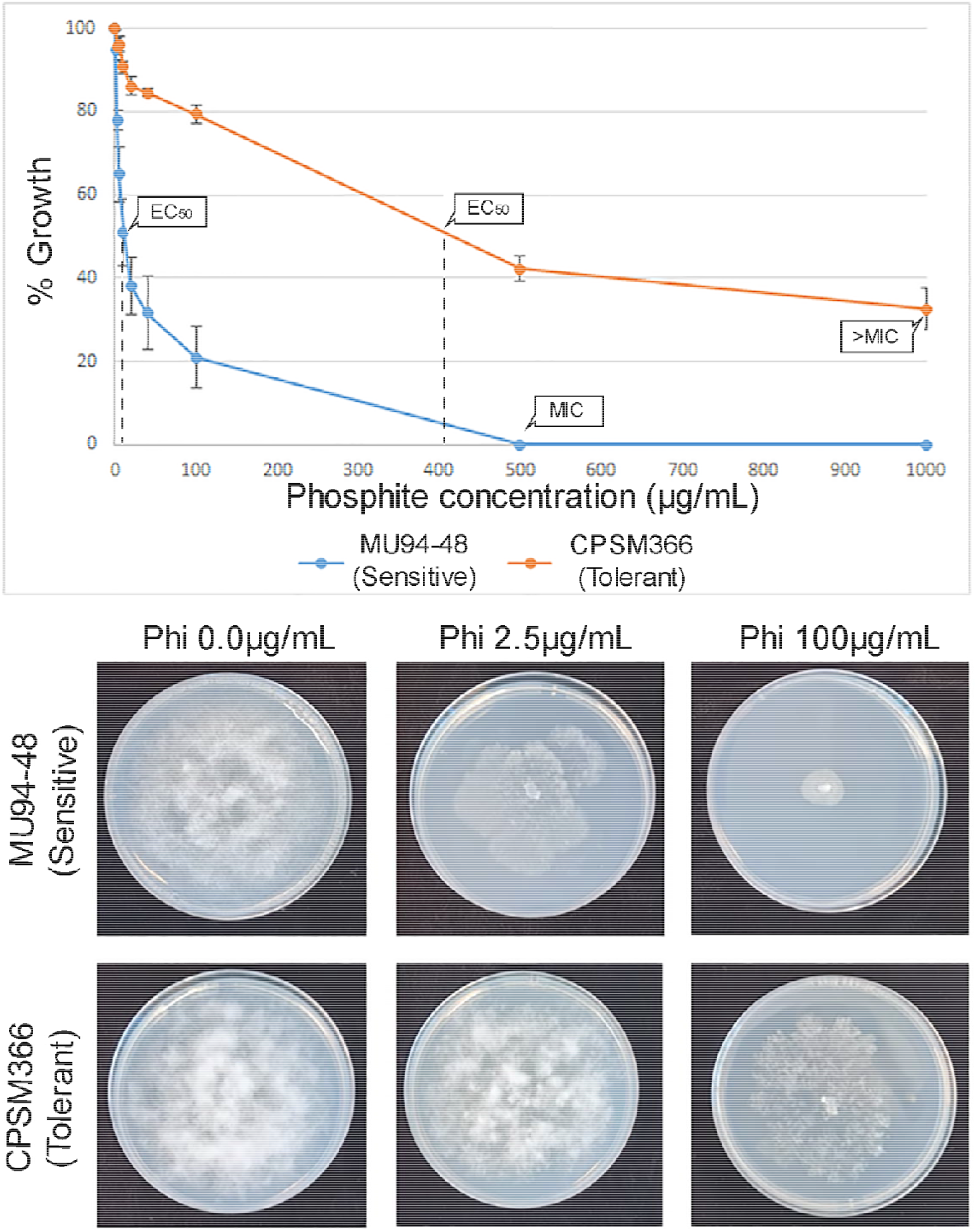
*In vitro* growth inhibition of MU94-48 and a CPSM366 in response to phosphite treatment indicating the MIC and EC_50_ for each isolate. Images of mycelial growth on the phosphite supplemented media are also displayed.

For proteomic analyses, 0.0 μg/ mL and 2.5 μg/ mL were used as these represent an untreated and sub-lethal dose of phosphite resulting in reduced growth of 20% and 4% for MU94-48 and CPSM366, respectively.

### 3.2 Phosphite induced significant alterations in the proteome of MU94-48 and CPSM366

To determine the biochemical differences between MU94-48 and CPSM366 treated with phosphite, the soluble intracellular proteome of each isolate was compared by qualitative and quantitative proteomic analyses. 1393 proteins were unique to MU94-48, 280 were unique to the CPSM366 and 1572 were common between the two isolates (Figure 2). Of the common proteins, 171 were higher in abundance in MU94-48 compared to CPSM699 and 90 were lower in abundance. 1311 were not differentially abundant. Overall, there was a significantly higher number of proteins identified in MU94-48, with 42.9% only observed in this isolate.

**Figure 2.**
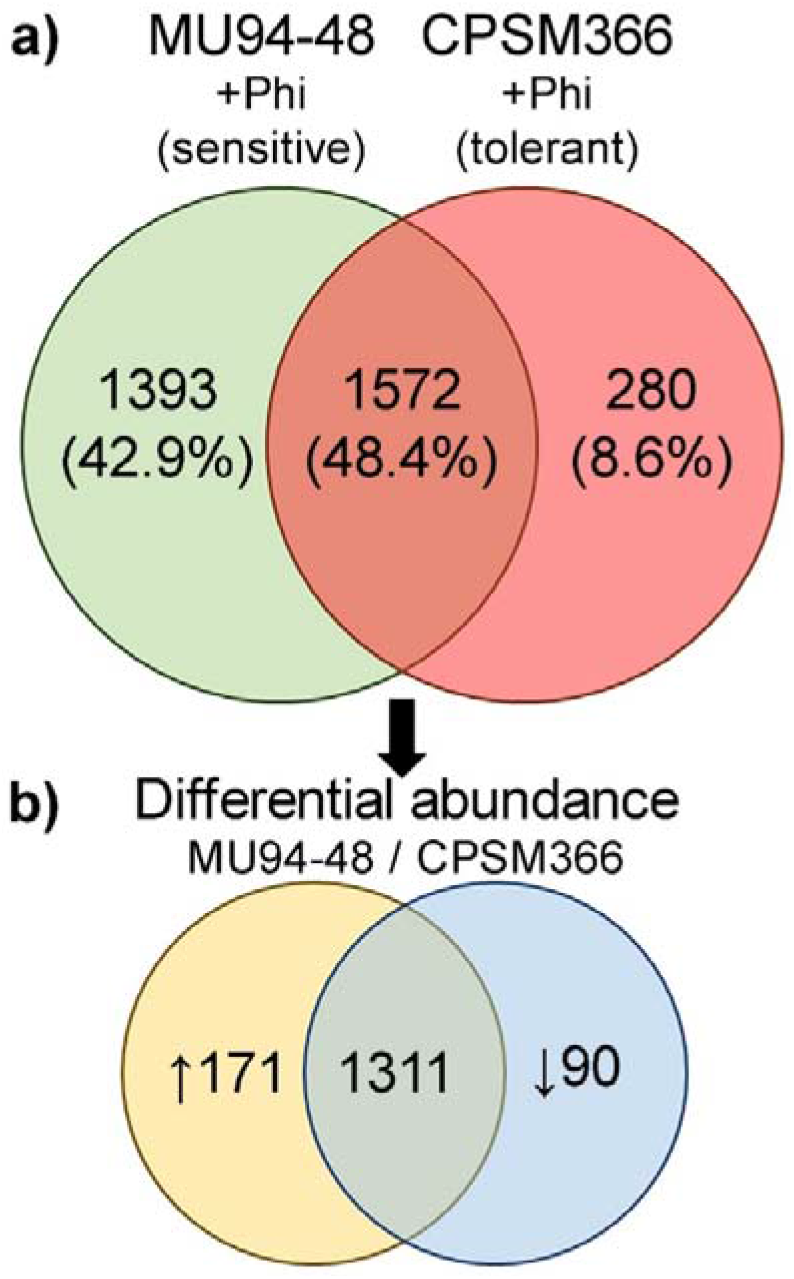
Protein identification in phosphite treated MU94-48 and CPSM366. **a)** Indicating the total number of unique and common proteins between the sensitive and tolerant isolates. **b)** Differential abundance of the common proteins by label-free quantification shows the increased and decreased abundance of proteins in phosphite-treated MU94-48 compared to phosphite-treated CPSM366. The number of proteins identified in all samples obtained for the *in vitro* assay are shown in Supplementary material 1.

### 3.3 Phosphite induces a stress response in MU94-48

GO enrichment analysis and differential protein abundance between the MU94-48 and CPSM366 were used to elucidate biochemical changes from phosphite treatment (Figure 3). In MU94-48, an enrichment of putative stress response proteins was observed including glutathione S-transferases (GST), thioredoxins, peptidases, proteasomes and proteolytic enzymes ^43^. These may be indicators of programmed cell death or attempts of the pathogen to stay alive. Putative stress proteins were not observed in CPSM366.

**Figure 3.**
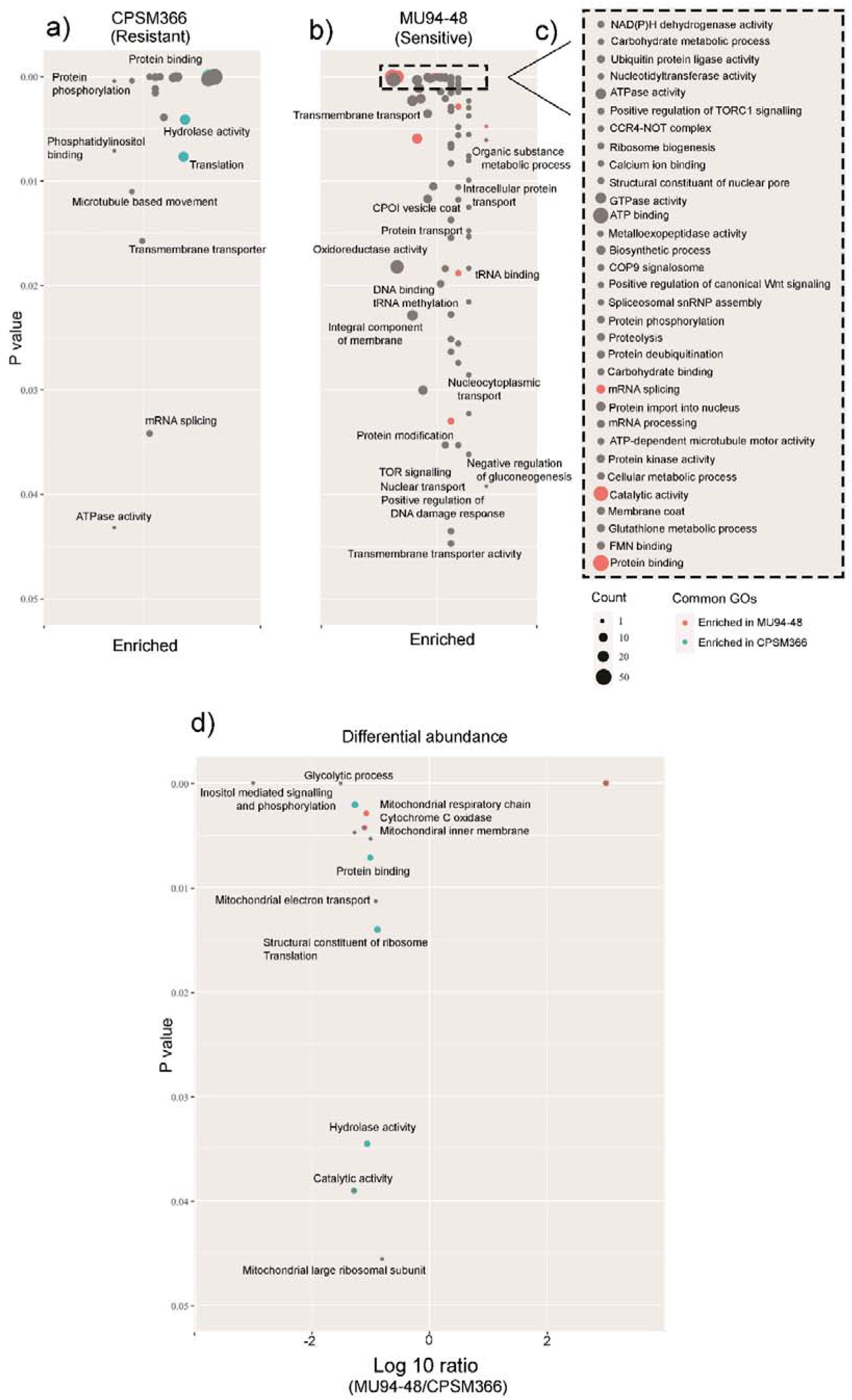
GO enrichment and differential abundance between MU94-48 and CPSM366 treated with phosphite. **a)** and **b)** show GO enrichment in the phosphite treated tolerant and sensitive isolates respectively, where the P value is generated by GO enrichment and each point is separated on a log10 scale generated by each GO count to the total GO count for the sample set. **d)** The enriched gene ontologies in the sensitive isolate at P < 0.001. **d)** Differential abundance of proteins between the sensitive and tolerant isolate. Ratios are generated by the peptide signal of each protein in MU94-48 compared to CPSM366. P values depict the significance of the differential abundance.

### 3.4 An increase in signalling in response to phosphite treatment

Enrichment of GOs that are associated with protein signalling including phosphorylation (eg. ATPases, GTPases, PKs), and ubiquitination were observed ^44^ (Figure 3). GOs that are associated with positive regulation of DNA damage were enriched in addition to tRNA binding, DNA binding, tRNA methylation and mRNA splicing and processing. Inositol signalling and phosphorylation were significantly lower in abundance in MU94-48 coinciding with the enrichment of phosphatidylinositol binding in CPSM366. Positive regulation of TORC1 signalling and Wnt transmembrane signalling GOs were enriched. This indicates stress response signalling, increase in gene expression to compensate for the loss of biological material, and signalling to coordinate cellular processes during stress conditions ^45,46^.

### 3.5 More transporters in MU94-48 than CPSM366 when treated with phopshite

A higher diversity of transporters was found in MU94-48, including transmembrane transporters, and intracellular transporters, along with facilitators of transport such as COPI vesicle coatomer proteins, COP9 signalosome, armadillo-like proteins and clathrin proteins (Figure 3). MU94-48 may be using transporters to pump out phosphite or toxic biproducts of phopshite metabolism as a means to facilitate the high level of signalling.

### 3.6 Phosphite alters mitochondrial respiration in MU94-48

A significantly lower abundance of mitochondrial-associated ontologies were observed in MU94-48 as indicated by mitochondrial respiratory chain, cytochrome C oxidase, mitochondrial electron transport and mitochondrial ribosomal subunit GOs. This could be a result of oxidative stress in MU94-48 or may suggest that similarly to other fungicides, phosphite targets mitochondrial respiration ^47,48^. The KEGG ontologies between phosphite-treated MU94-48 and CPSM366 did not show changes in disctinct metabolic pathway clusters but rather showed general metabolic disorder (Supplementary material 2). However, the disordered KEGG orthologues in MU94-48 suggest that phosphite is exerting a cytotoxic effect on the sensitive isolate.

We then examined the proteome of MU94-48 and CPSM66 compared to their respective untreated controls to ensure that these observations were not artifacts of the isolates. It was observed that the proteome profile of untreated and phosphite-treated MU94-48 and CPSM366 are comparable to those found between MU94-48 (Supplementary material 3). This includes the enrichment in putative stress proteins such as oxidoreductase activity, intracellular signalling and protein activation, and gene expression-related ontologies. Additionally, a significant reduction in inositol biosynthesis and signalling, mitochondrial electron transport-related GOs were observed, reflecting the observations between the two phosphite-treated isolates. Similarly, we compared the proteome of untreated and phosphite-treated CPSM366 and found less biochemical responses than when MU94-48 was compared to CPSM366. The full GOs, KEGG orthologues and gene functions for all *in vitro* qualitative and quantitative comparisons are shown in Supplementary Material 4.

### 3.7 Differential abundance of proteins between untreated and phosphite treated *Lupinus angustifolius*

We hypothesise that phosphite in the plant physiology by facilitating plant immunity to prevent infection by phytopathogens such as *P. cinnamomi*. To investigate this, the proteome of *L. angustifolius* treated with and without phosphite were compared. 498 proteins were unique in the untreated Lupin sample and 1725 were unique in the phosphite-treated Lupin (Figure 4). 245 proteins were common between the untreated and treated lupin samples of which 46 were significantly higher in abundance when phosphite was applied and 4 were significantly lower.

**Figure 4.**
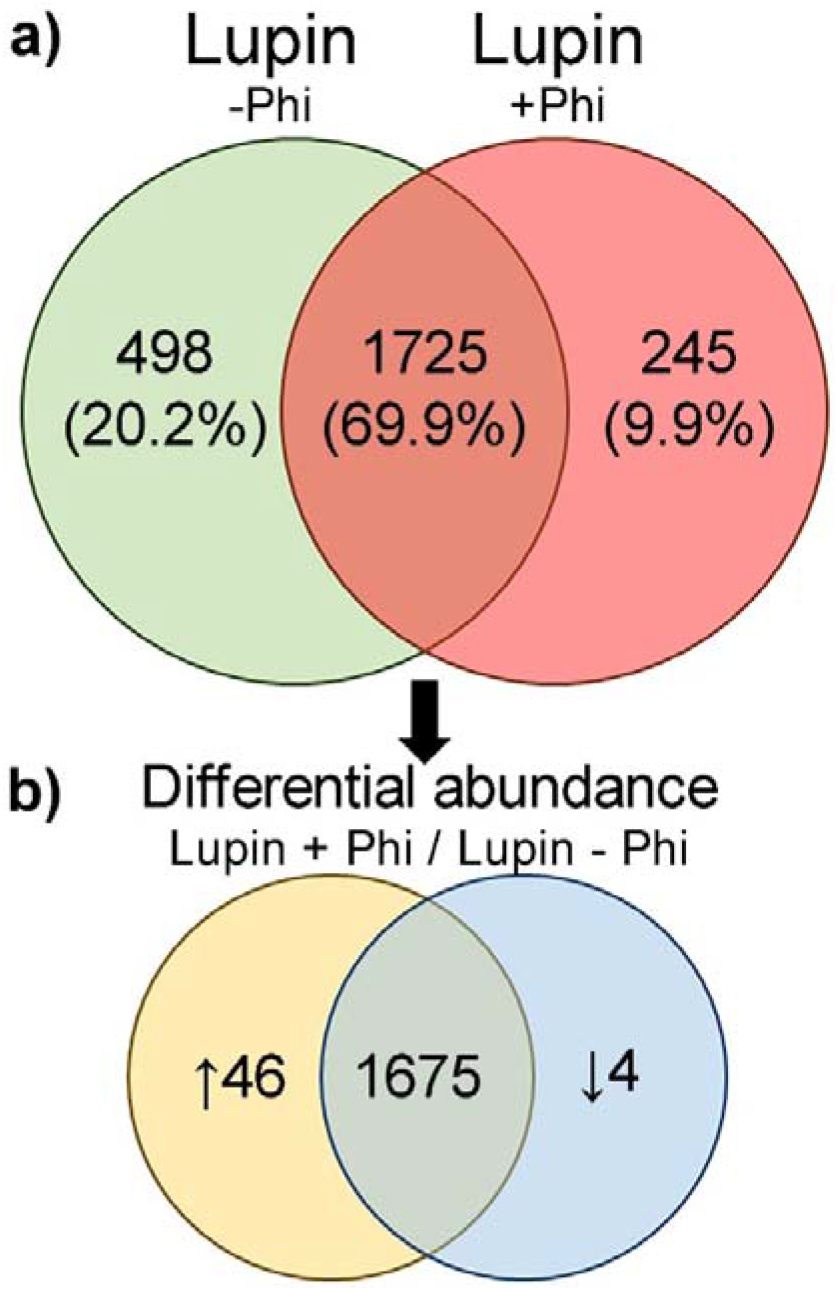
Protein identification between untreated and phosphite treated lupins. **a)** Indicating the total number of unique and common proteins between the untreated and phosphite-treated lupin. **b)** Differential abundance of the common proteins by label-free quantification shows the increased and decreased abundance of proteins in phosphite-treated lupin compared to the untreated control. The number of proteins identified in all samples obtained for the *in planta* assay are shown in Supplementary material 1.

### 3.8 Phosphite increased the abundance of photosynthetic and carbon fixation proteins in *L. angustifolius*

GO enrichment and KEGG pathway enrichment analyses were used to examine biochemical effects in lupin treated with phosphite, (Figure 5 and 6). It was observed that GOs containing proteins that are associated with photosynthesis and starch metabolism were significantly enriched in phosphite-treated lupins. Photosynthesis, photosystem units, geranylgeranyl reductase, chlorophyll biosynthesis, carbon fixation and TCA cycle were all enriched in phosphite-treated lupins. Clustering of KEGG ontologies only found in the phosphite-treated lupin mapped to photorespiration, photosynthesis, reductive pentose phosphate cycle, isoprenoid biosynthesis and carbon fixation (Figure 6). As lupin seedlings were grown under artificial light with roots exposed leading to the development of photosynthetic-capable tissues, these were excised as part of the excised root tip for protein extraction (Figure 7).

**Figure 5.**
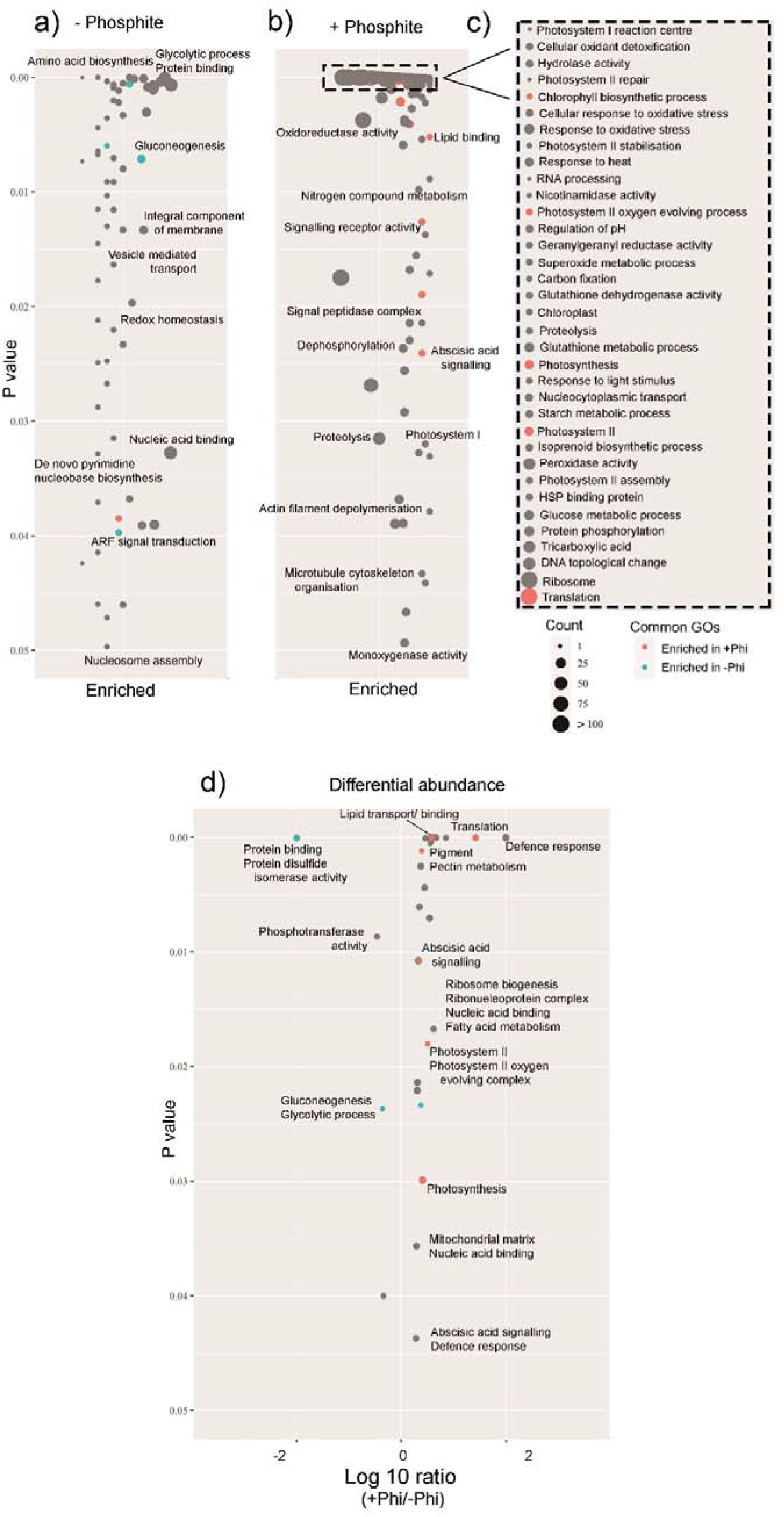
GO enrichment and differential abundance between untreated and phosphite-treated lupin. **a)** and **b)** show GO enrichment in the untreated and phosphite-treated lupin respectively, where the P value is generated by GO enrichment and each point is separated on a log10 scale generated by each GO count to the total GO count for the sample set. **c)** The enriched gene ontologies in the phosphite-treated lupin at P < 0.001. **d)** differential abundance of proteins between the untreated and treated lupin. Ratios are generated by the peptide signal of treated lupin compared to the untreated control. P values depict the significance of the differential abundance.

**Figure 6.**
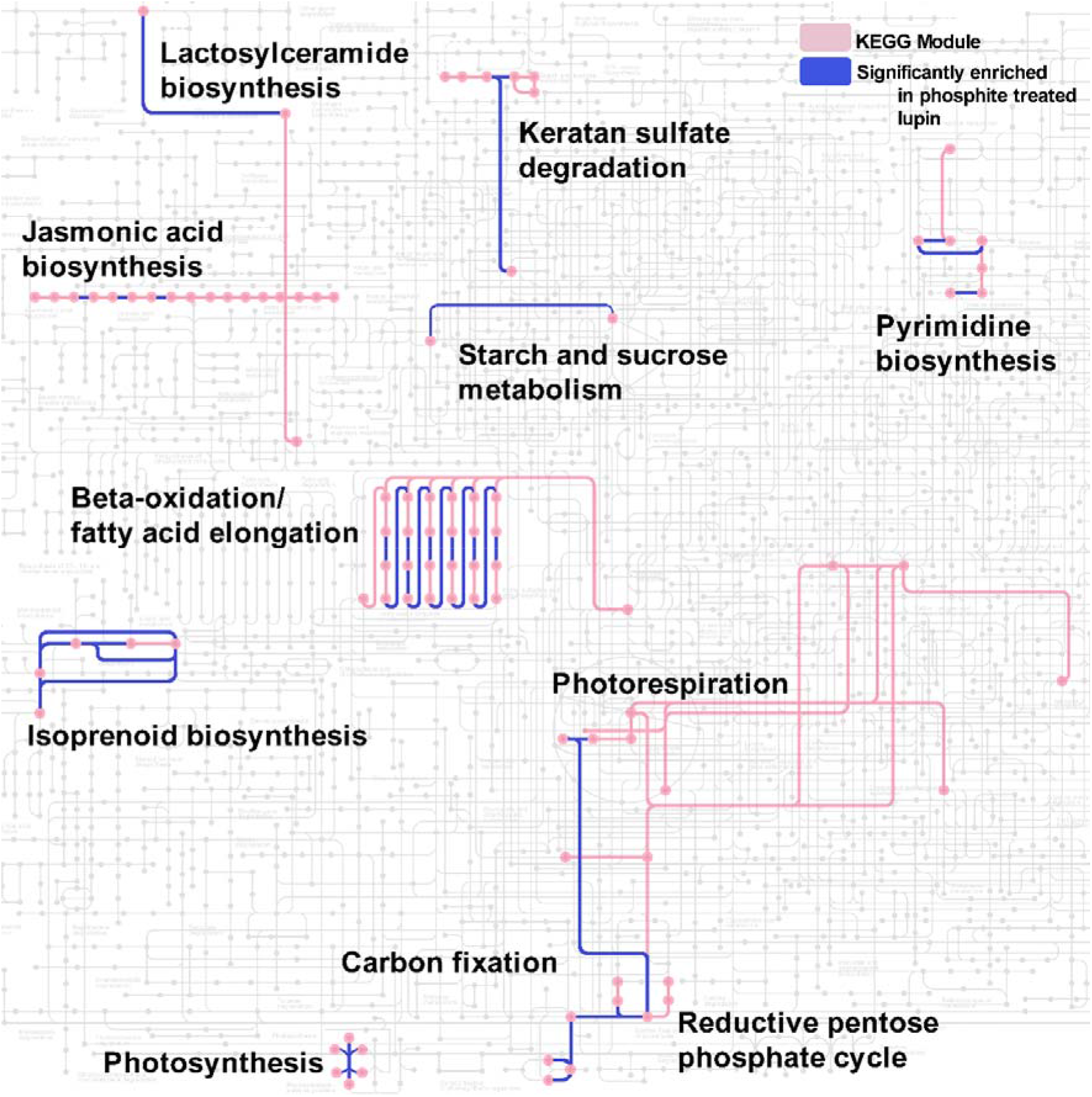
A representative set of the KEGG pathways enriched in the phosphite-treated lupin. Each dot represents a KEGG entry, and each line represents individual KEGG ontology identifiers. Pathway sets from overrepresented modules are labelled.

**Figure 7.**
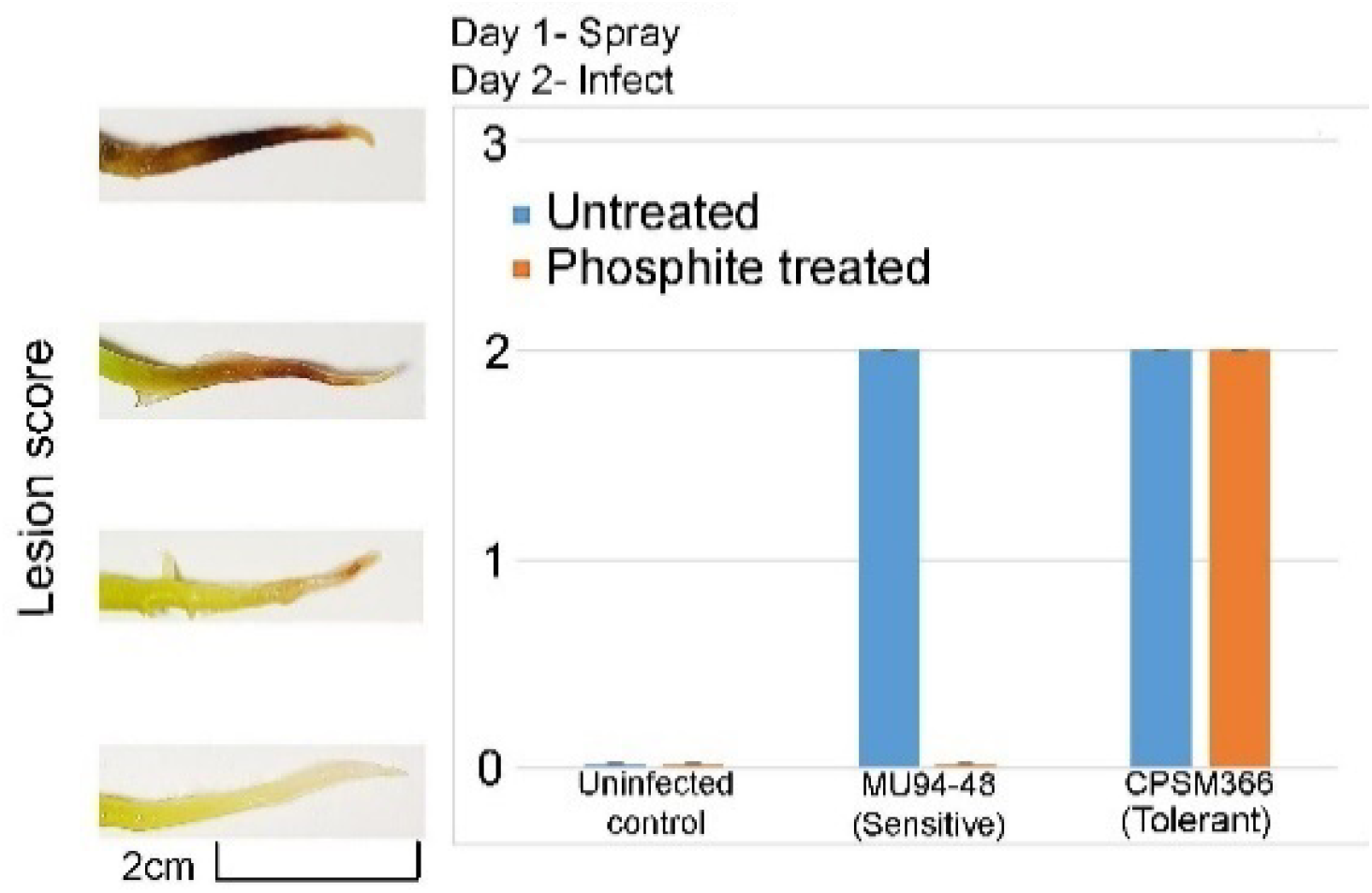
Lesion scores of lupin roots when treated with phosphite and infected with the two isolates of *P. cinnamomi*. Lesion scores were taken on day 3.

Proteins associated with glucose, starch, and lipid metabolism GOs were enriched in the phosphite-treated lupin along with related KEGGs such as starch and sucrose metabolism, beta-oxidation and fatty elongation and lactosylceramide. Gluconeogenesis was reduced in abundance which supports the utilisation of carbohydrates and sugars as an energy source to fuel the heightened metabolism. Similarly, KEGG orthologues that are associated with lipid metabolism were also observed as indicated by the clustering of identified KEGG pathways only found in the phosphite-treated lupin (Figure 6). Phosphite treatment results in increased abundance of proteins that are associated with photosynthesis, carbon metabolism and energy production in lupin.

### 3.9 Transcriptional activities in phosphite-treated *L. angustifolius* are enhanced

As a complement to the abundance of photosynthetic and metabolic gene ontologies, gene expression was also overrepresented. Translation, ribosomal proteins, DNA topological change, and RNA processing were all enriched in the phosphite treatment (Figure 5). This is also shown by the clustering of KEGG pathways in the phosphite-treated lupin including pyrimidine biosynthesis and keratan sulfate degradation (Figure 6). Hence, an increase in gene expression and biosynthesis accompanies massive physiological and metabolic changes.

### 3.10 Phosphite triggers the accumulation of defence-related proteins in lupin

Phosphite has previously been reported to increase the production of defence-related molecules in plants, particularly salicylic and jasmonic acid ^7,8,49^. An enrichment of defence-related GOs was observed in the phosphite-treated lupin. These included proteolysis, oxidoreductase activity, hydrolases, cellular oxidant detoxification and response to oxidative stress (Figure 5). Secretory peroxidases, superoxide metabolism, abscisic acid signalling, and defence response GOs encompassing genes such as steroid chaperones, programmed cell death and apoptosis were also enriched. Several secondary metabolites were enriched including nicotinamidase and isochorismatase were also found only in the phosphite-treated lupin.

Precursors to the salicylic acid (SA) pathway including isochorismatase and actin depolymerisation GOs were enriched in the phosphite-treated lupin. KEGG analysis revealed components associated with the jasmonic acid (JA) pathway were only found in the phosphite-treated lupin. This shows that more defence-related proteins including secondary metabolites were enriched and higher in abundance in the phosphite-treated lupin.

### 3.11 The effect of phosphite on MU94-48 and CPSM366 isolates during host infection

To test the effect of phosphite during host infection*, L. angustifolius* was treated with phosphite and subsequently inoculated with the sensitive or tolerant isolate of *P. cinnamomi* (Figure 7). Untreated lupin infected with MU94-48 and CPSM366 developed comparable lesions. When phosphite-treated *L. angustifolius* was infected with MU94-38 no lesion was observed showing sensitivity to phosphite *in planta*. When inoculated with CPSM366, there was no reduction in lesion score compared to the untreated lupin.

## 4 DISCUSSION

The mode of action of phosphite in *Pythophthora* pathosystems is a central question in the development of future management strategies. As resistance to the only effective chemical for the control of *P. cinnamomi* and other ooomycetes emerges, the pressure to find resistance genes or alternative management strategies mounts. An understanding of the mode of action of phosphite could aid in the development of better oomyceticides, which in turn would contribute to improving existing management strategies. Previous literature has suggested both direct and indirect mechanisms of phosphite on the pathogen and its host plants however the biochemical mechanisms in which these occur have not been defined. Our study aims to deconvolute the biochemistry of this system and gain a clearer insight into the pathways altered by phosphite. *L. angustifolius* (lupin) is used as a model as it is susceptible to *P. cinnamomi* infection, it has a published genome sequence required for proteomics work, and it grows rapidly compared to native and horticultural *P. cinnamomi* hosts ^39,50^.

The *in vitro* growth assay demonstrates that phosphite has a direct inhibitory effect on mycelial growth in *P. cinnamomi*. The EC_50_ of previously reported phosphite-sensitive *P. cinnamomi* isolates ranges between 4 μg mL^−1^ and 25 μg mL^−1 20,51^. The EC_50_ MU94-48 falls within this range and can be considered highly sensitive to phosphite. Reported tolerant isolates of *P. cinnamomi* have EC_50_ concentrations up to 150 μg mL^−1^ and other species of *Phytophthora* have EC_50_ values up to 350 μg mL^−1^, of which CMSP366 exceeds ^20,51–53^. *P. citrophthor* and *P. syringae* isolates screened for sensitivity to phosphite were considered sensitive when their EC_50_ values were below 25 μg mL^−1^), whereas *P. nicotianae* isolates with EC_50_ values above 75 μg mL^−1^ were considered moderately resistant or resistant ^24^. CPSM366 is therefore considered as highly resistant to phosphite, highlighting the need for understanding its biochemistry and the development of alternative management strategies.

A proteomic approach was taken to understand how phosphite interacts with the two *P. cinnamomi* isolates MU94-48 and CPSM366 and causes growth reduction. The data obtained from this work was used to build a model suggesting the possible mechanisms in which phosphite directly affects the pathogen (Figure 8). In the presence of phosphite, an abundance of putative stress proteins is observed in the sensitive isolate (Figure 8a). The pathogen is trying to find ways to cope with the influx of phosphite by expressing GSTs and thioredoxins ^54^. Hydrolase, proteolysis and peptidase proteins could also be produced by the pathogen in this case for nutrient recycling, detoxification of xenobiotics, or could be products of cell death caused by phosphite ^43,55^. Putative stress response proteins were only observed in MU94-48 indicating that CPSM366 does not undergo abiotic stress when treated with phosphite.

**Figure 8.**
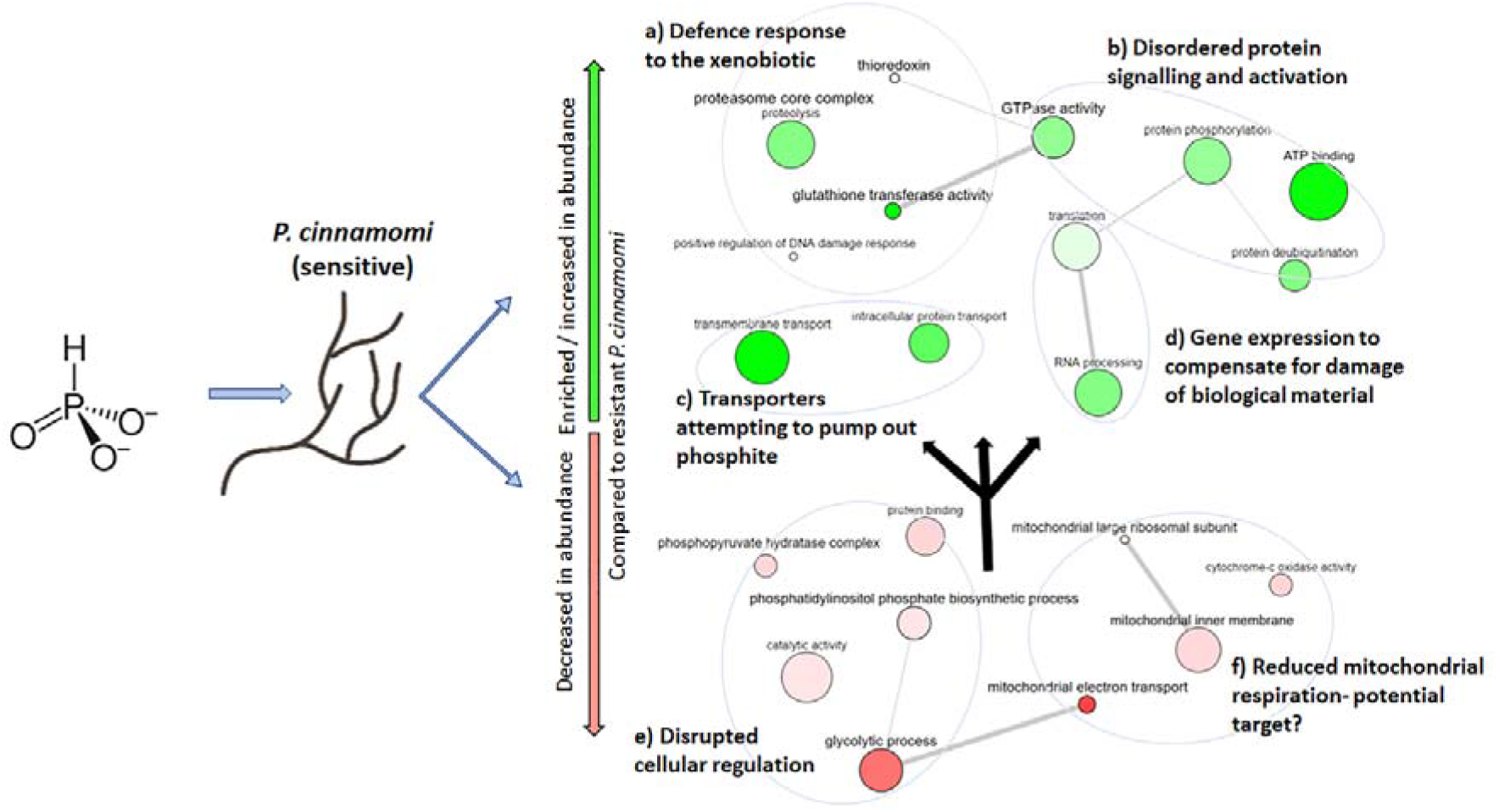
A proposed model on the direct effect of phosphite on *P. cinnamomi* MU94-48 from the combination of qualitative and quantitative comparisons between MU94-48 and CPSM366. Black arrows indicate a possible cascading effect of core biochemical pathways that are disrupted as a result of phosphite treatment. The size of bubbles represents enrichment relative to each other for visualisation.

Signalling molecules related to stress response were also induced along with the disruption of the regulation of cellular processes caused by the exposure of MU94-48 to phosphite (Figure 8b). Several core regulators of cell growth, metabolism and signalling were altered as a result of phosphite treatment. For example, TORCI signalling was enriched in MU94-48, which is involved in aspects of cell growth and metabolism ^46^. This indicates that the pathogen attempts to cope with disrupted cell growth and metabolism caused by phosphite. Positive response to regulation of DNA damage and increase in gene expression-related ontologies shows that phosphite could cause DNA damage and subsequent increase in gene expression to compensate for the loss of DNA, proteins and cell material through biosynthesis (Figure 8d) ^56^. Canonical Wnt proteins are core regulators of development, another biochemical pathway that is altered by phosphite. Inositol signalling and phosphorylation, which play important roles in aspects of cell growth, structure and signalling, were significantly lower in abundance in MU94-48 (Figure 8e). These mechanisms of coping with extensive damage do not seem to be effective enough to protect the sensitive isolate from damage by phosphite.

Greater diversity of membrane transporters were observed in the proteome of MU94-48 than in CPSM366 (Figure 8c). These transporters include ABC and MSF-type, which have been described to participate in the removal of xenobiotics from the cell ^57,58^. Additionally, facilitators of intracellular transport were also enriched in MU94-48 suggesting that, unlike CPSM366, MU94-48 attempts to remove the toxic xenobiotic.

Compared to the tolerant isolate CPSM366, several constituents of the mitochondrial respiratory pathway were reduced in abundance in the sensitive isolate (Figure 8f). In fungal pathogens, programmed cell death can be triggered by mitochondrial-initiated signalling that can be activated by cell damage, exposure to toxic xenobiotics and oxidative stress ^59^. During these stress conditions, reactive oxygen species can disrupt components of the mitochondrial respiration chain causing subsequent cell death ^60–62^. As this is a core pathway for cellular function, mitochondrial respiration is commonly used as a target for chemical control of fungal phytopathogens by single-site fungicides ^48,63^. For example, strobilurins are a broad-spectrum class of fungicide applied to control fungal crop diseases. Strobilurin binds cytochrome b complex III, inhibiting mitochondrial respiration ^64^. Similarly, azoxystrobin blocks electron transport in mitochondrial respiration by blocking electron transport ^64,65,66^. Evidence presented here might suggest a broader inhibitory activity in the case of phosphite as it seems to be affectiving the pathogen’s metabolism at multiple levels. Many fungicides have multi-site modes of action, where several biochemical processes are disrupted such as chlorothalonil, folpet, thiram, sulphur and copper ^67,68^.

The metabolic, signalling, regulatory and stress responses were not observed in CPSM366. If phosphite acts as a multi-site oomyceticide, CPSM366 has likely developed physiological adaptation to tolerate phosphite or alteration in target sites ^69,70^.

It has been suggested that phosphite alters the plant system to better cope with potential attacks from oomycete pathogens by priming the plant immune system ^7,8^. In addition, phosphite has been reported to act as a biostimulant in plants ^71^. A proteomic approach was used to obtain a detailed biochemical snapshot of the proteins differentially abundant in phosphite-treated *L. angustifolius*. The proteomic data obtained from the *in planta* assay was used to generate a model that describes how phosphite alters plant metabolism and induces an increase in defence-related proteins (Figure 9).

**Figure 9.**
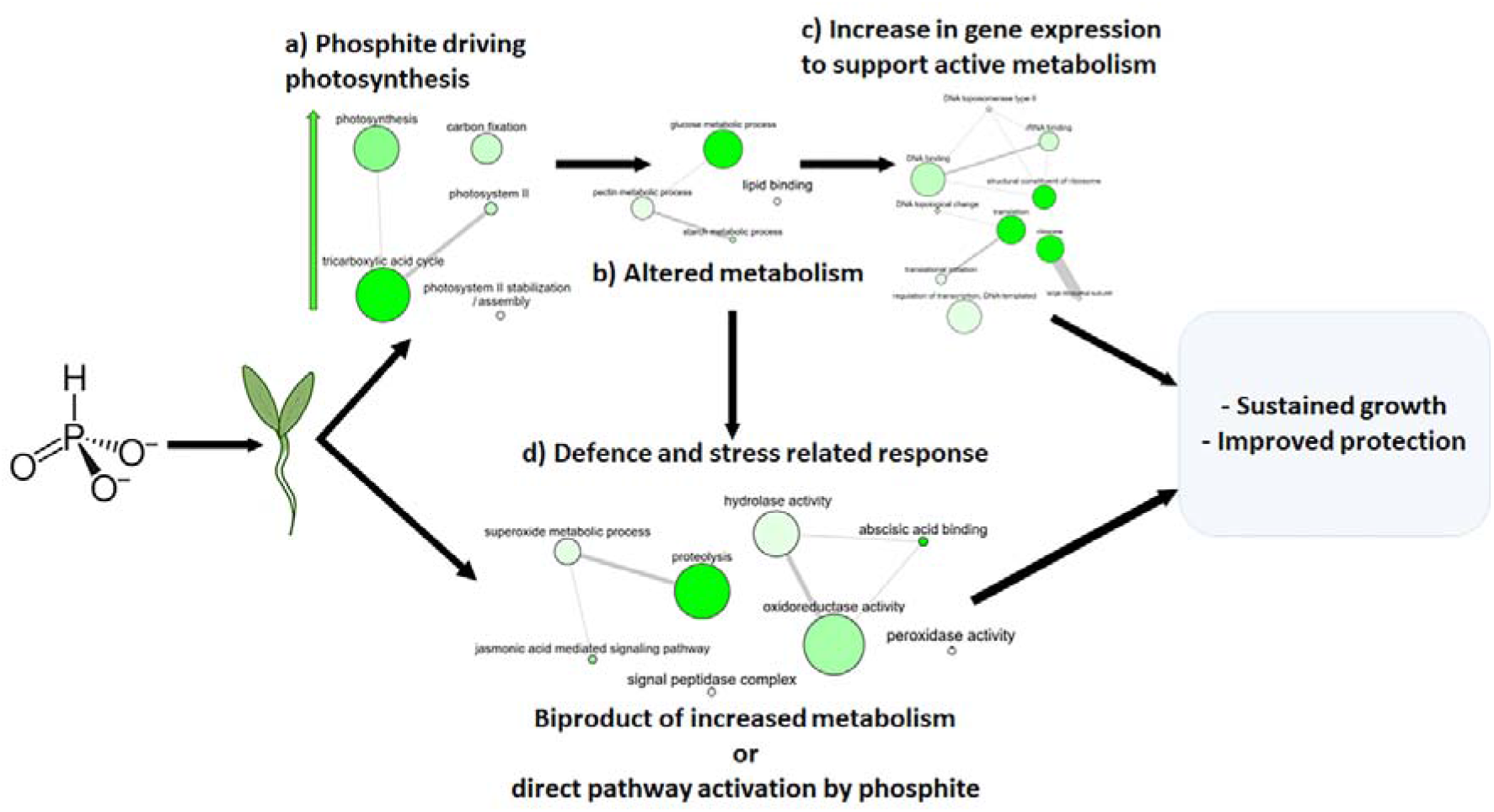
A proposed model on the effects of phosphite on *L. angustifolius* from the combination of qualitative and quantitative proteome data between the untreated and treated lupin.

The biochemical differentiation between phosphite treated and untreated *L. angustifolius* suggests an increase in metabolic activity. Constituents of the photosynthetic process, carbon fixation and citric acid cycle suggest that phosphite is driving metabolism in plants (Figure 9a). In addition, an increase in lipid and carbohydrate metabolism was observed, indicating that stored carbohydrates can be metabolised for energy generation (Figure 9b). The reductive pentose phosphate cycle was observed in the KEGG ontologies of the phosphite-treated lupin and generates NADPH that feeds back into glycolysis ^72^. Gluconeogenesis is significantly reduced in the phosphite-treated lupin highlighting that sugars are utilised for energy production in this state, not stored. This is reinforced by enrichment in gene expression to keep up with the metabolism and to cope with the increase in photosynthesis through the process of biosynthesis (Figure 9c).

An increased metabolism has been observed in other phosphite-treated plants compared to untreated controls along with its biostimulant effect ^22,71,73,74^. The application of phosphite improves yield, biomass, fruiting and growth, with a field trial applying phosphite in avocado plantations showing a significant increase in fruit production ^75,76^. In potato, phosphite application resulted in reduced seedling emergence time, increased leaf size and biomass ^77^. This ‘greening effect’ has been demonstrated amongst other fungicides in agriculture, where yield, biomass, leaf surface area, and protein content are increased as a result of fungicide application ^78–80^. The response observed in the phosphite-treated lupin seedlings is similar to the greening effect, where phosphite boosts the metabolism of the plant. Non-leaf plant structures such as roots, stems, flowers and seeds have photosynthetic potential when exposed to light and carbon fixation has been reported to occur in many plant roots along with green roots ^81–83^.

Defence-related gene expression has been demonstrated in several phosphite-treated crops. In potato, phosphite treatment caused a significant increase in transcription of salicylic and jasmonic acid ^8^. Proteomic analysis of a similar system showed an increase in the abundance of peroxidases, glutathione S-transferase and proteinase inhibitors ^84^. In the present study, enrichment of defence and stress-related proteins in *L. angustifolius* were detected as a result of phosphite treatment, which may act in favour of the defence response of the plant.

Secretory peroxidases were also enriched in the phosphite-treated lupin which can oxidise toxic compounds and have functional roles in defence and biosynthesis ^85^. Superoxide metabolism and phospholipid binding composed of annexin genes were also enriched. Superoxide dismutase is used in plant defence against reactive oxygen species ^86,87^. Isoprenoids identified in the phosphite-treated lupin are not only carriers in photosynthetic and respiratory electron transport and have additional functions as antioxidants ^88^. Defence-related ontologies with gene functions related to steroid chaperoning indicating programmed cell death and apoptosis were also abundant ^89^. Signalling and binding of abscisic acid, a key hormone involved in signalling during stress and defence response to abiotic and pathogens was significantly higher in abundance in the phosphite-treated lupin. Nicotinamidase and isochorismatase are involved in plant growth, hydrolase activity and synthesis of salicylic acid ^90–93^ JA pathway is associated with defence response against necrotrophic microbial pathogens and abiotic stresses and the SA pathway is involved with biotic stressors, cell death and hypersensitive responses in plants ^94,95^.

In this system, phosphite is driving both metabolism and defence by directly promoting their biochemical pathways ^8^. The production of stress molecules such as abscisic acid can at the same time be used as signalling molecules to trigger plant immunity ^96^. The production of defence and stress-related molecules in the plant could also be acting in response to phosphite as a xenobiotic substance ^97,98^. In the field, the ideal use of phosphite part of a preventative strategy for the management of *P. cinnamomi* infection ^11–13^. The *in planta* assay demonstrates that phosphite application does not reduce the observed lesion when lupin plants are infected with the tolerant isolate. If the elevated defence response impacts the colonisation ability of *P. cinnamomi*, it is not observed in this system. Potentially the induction of defense molecules by phosphite in this system may not have reached a sufficient amplitude for host resistance.

## 5 CONCLUSION

Our data present a new perspective on the mode of action of phosphite. We have provided evidence to propose models of the direct and indirect mode of action of phosphite. This data demonstrates a comprehensive snapshot of the metabolic dysregulation in the sensitive isolate when treated with phosphite, suggesting it is a substrate for one or more core biochemical pathways. The tolerant isolate is likely to have adapted to block phosphite from entering the cell, or if phosphite targets constituents of mitochondrial respiration, adapted to block this interaction. Our model proposes an alternative avenue of plant responses to phosphite, where phosphite drives multiple biochemical pathways and as a bi-product, more defence-related proteins are produced. We proposed probable mechanisms based on proteomic data that indicates a bi-modal mode of action of phosphite on both the pathogen and the host plant. Further studies are required to determine systemic changes in phosphite-treated *L. angustifolius* such as changes in photosynthesis and respiration in leaves to confirm the current observations. Additionally, the accumulation of phosphite in the plant tissue and subsequent uptake by the pathogen cannot be excluded. The outcome of this study presented opportunities for functional validation to determine the phosphite mode of action in crop protection.

## Availability of data and materials

Spectral data used for this study are available at Figshare (DOI: 10.6084/m9.figshare.20026214).

## Acknowledgements

Proteomics International provided funding for the project. Curtin University provided funding for sample preparation through the postgraduate maintenance fund. KCT, FLR and SJ are supported by the Centre for Crop and Disease Management, a joint initiative of Curtin University and the Grains Research and Development Corporation (CUR00023). We thank Dr Lars Kamphuis from the Centre for Crop and Disease Management and CSIRO for providing the seed stocks. We also thank Johannes Debler for his technical input for functional annotation. We thank Prof. Giles Hardy and Dr Bill Dunstan from the Centre of Phytophthora and Science Management at Murdoch University for providing the *P. cinnamomi* isolates. We also thank Mr Leon Lenzo for his critical feedback on the manuscript and Dr Paula Moolhuijzen for her input into bioinformatic visualisation.

**Supplementary material 1-** Protein identification of *in vitro phosphite* treated MU94-48 and CPSM366 treated with phosphite and of the phosphite treated lupin roots. FDR was set to <1%.

**Supplementary material 2-** Th KEGG pathways identified from the protenis only found in MU94-48. Green lines indicate the KEGG pathways assigned to the unique proteins in phopshite treated MU94-48 compared to the untreated control. There are no clear clusters of pathways in this sample indicating that phosphite affects multiple processes in the sensitive isolate.

**Supplementary material 3-** Representative GOs obtained from phosphite treated MU94-48 and CPSM366 compared to their respective untreated controls.. **a)** Enriched GOs in the proteins uniquely identidied in untreated MU94-48 compared to treated MU94-48. **b)** GOs associates with the differentially abundant proteins between untreated and phopshite treated MU94-48. **c)** Enriched GOs in the proteins uniquely identidied in treated MU94-48 compared to tnreated MU94-48. **d)** Enriched GOs in the proteins uniquely identidied in untreated CPSM366 compared to treated CPSM366. **e)** GOs associates with the differentially abundant proteins between untreated and phopshite treated CPSM366. **f)** Enriched GOs in the proteins uniquely identidied in treated MU94-48 compared to tnreated MU94-48. Green bubbles in **a), c), d)** and **f)** indicate significantly enriched GOs, green bubbles in **b)** and **e)** indicated significantly higher in abundance in the treated samples and red bubbles indicate significantly lower abundance in the treated samples compared to untreated samples.

**Supplementary material 4-** The full list of GOs, Interpro and KEGGs for the qualitative and quantitative *in vitro* data set.

**Supplementary material 5-** The full list of GOs, Interpro and KEGGs for the qualitative and quantitative *in planta* data set.

