## Supplementary material 1 for "Proteomic analysis revealed that the oomyceticide phosphite exhibits multi-modal action in an oomycete pathosystem"

**Supplementary Table 1**

|  | | **Phosphite treatment** | **Total Spectra** | **PSM** | **Protein identifications >1 peptide** | **Protein identifications >2 peptides** |
| --- | --- | --- | --- | --- | --- | --- |
| ***In vitro*** | **MU94-48 (Sensitive)** | **-Phi** | 114369 | 45039 | 2038 | 1256 |
|  |  | **+Phi** | 142888 | 93885 | 2965 | 2181 |
|  | **CPSM366 (Tolerant)** | **-Phi** | 48942 | 14672 | 1343 | 746 |
|  |  | **+Phi** | 105305 | 38080 | 1852 | 1193 |
| ***In planta* treatment** | **-** | -Phi | 92622 | 57675 | 2058 | 1443 |
|  |  | +Phi | 88073 | 52220 | 1892 | 1334 |
