## Supplementary figures and images for "Proteomic analysis revealed that the oomyceticide phosphite exhibits multi-modal action in an oomycete pathosystem"

### Supplementary material 2

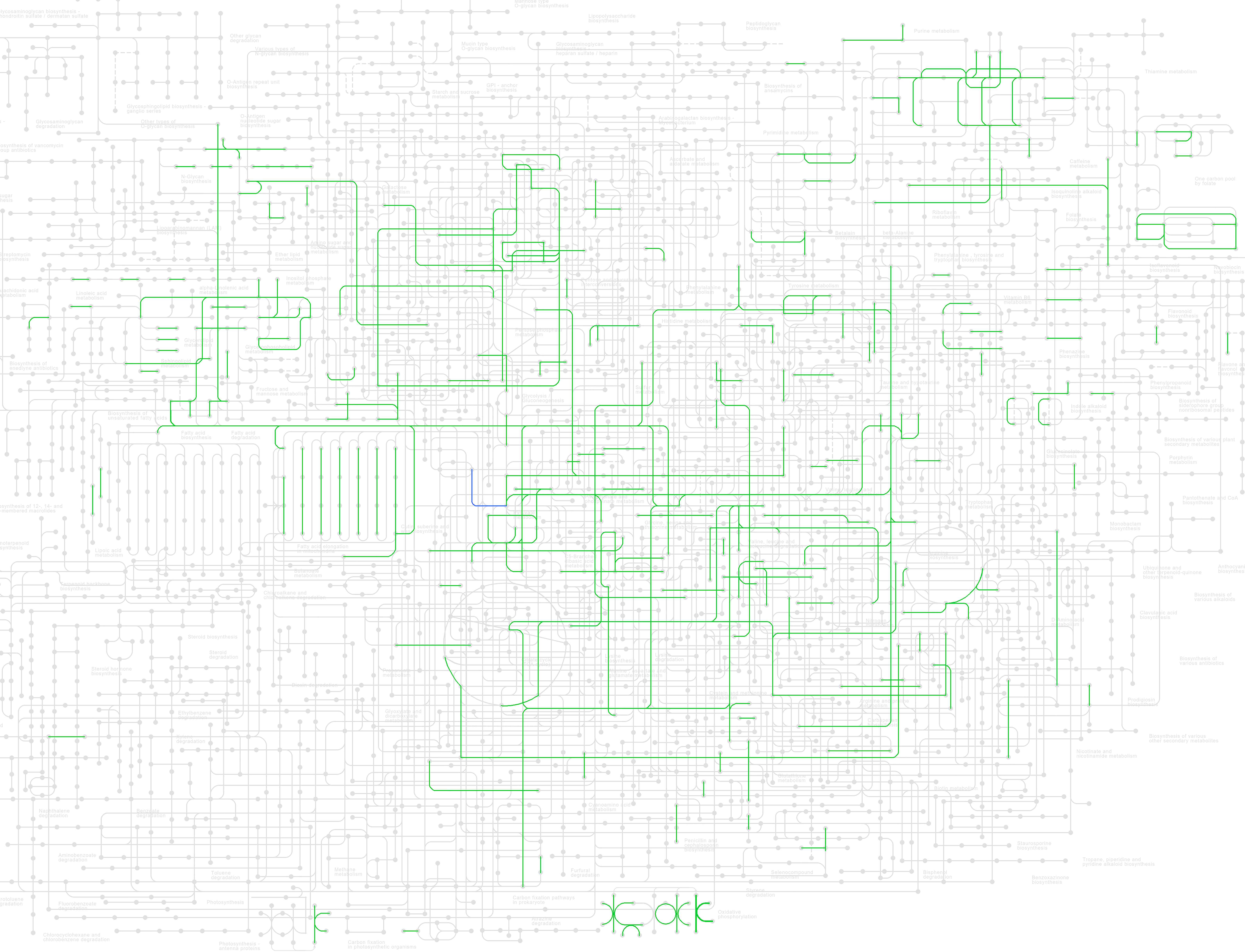
